# Consistent Effects of Pesticides on Community Structure and Ecosystem Function in Freshwater Systems

**DOI:** 10.1101/2020.02.03.932442

**Authors:** Samantha L. Rumschlag, Michael B. Mahon, Jason T. Hoverman, Thomas R. Raffel, Hunter J. Carrick, Peter J. Hudson, Jason R. Rohr

## Abstract

Predicting ecological effects of contaminants remains challenging because of the sheer number of chemicals and their ambiguous role in biodiversity-ecosystem function relationships. We evaluated responses of pond ecosystems to standardized concentrations of 12 pesticides, nested in four pesticide classes and two pesticide types. We show consistent effects of herbicides and insecticides on ecosystem function, but slightly less consistent effects on community composition. Effects of pesticides on ecosystem functions were often mediated by changes to biodiversity, and our analyses show that consistency in effects of pesticide types on functions was driven by functional redundancy among species. These results suggest that risk assessment of the thousands of registered chemicals on ecosystem responses could be simplified to a smaller number of chemical groups and to groups of functionally redundant taxa.

---

Freshwater systems are the most biodiverse in the world and provide important ecosystem services (*1*), yet they are imperiled by pesticide contamination (*2*). Two major challenges, among many, impede prediction of responses of freshwater ecosystems to pesticides. First, the extent to which individual pesticides have consistent effects on ecosystem functions and biodiversity is unknown. In the U.S. and Europe, tens of thousands of synthetic chemicals are registered, and in the U.S. >350 pesticides are applied annually (*3*). If the effects of pesticides are consistent within ‘pesticide classes’ (those with similar chemical structures) or ‘pesticide types’ (those targeting similar pests), then the complexity in predicting impacts of pesticides could be markedly reduced (*4*). Such consistency would improve efficiency of risk assessment and allow a greater focus on exceptions to general patterns. Second, the role of pesticides in biodiversity-ecosystem function relationships has not been elucidated (*5*). Historically, random and direct manipulations of single-trophic level communities and measurement of associated ecosystem processes (*6*) have established causality between biodiversity and ecosystem function (*7*). However, this approach overlooks the importance of anthropogenic factors, whose influences on communities are far from random (*8*), alter multiple trophic levels (*9*), and occur via direct and indirect pathways (*5*).

In an effort to suggest improvements to risk assessment, the objectives of the current study were to: 1) evaluate the consistency of effects across pesticide types, classes, and individual pesticides on ecosystem processes and communities, 2) assess whether the effects of pesticides on ecosystem processes and communities were the result of sublethal, non-target effects or changes in abundance of ‘targeted taxa’, and 3) determine if disruptions in ecosystem processes from pesticides were mediated by changes in biodiversity. We propose three hypotheses. First, ecosystem processes respond consistently to different pesticides within pesticide types because taxonomically related community members often have similar functional roles within the ecosystem. So, reductions in the abundance of taxa of a single group (e.g., green algae) might be specific to an individual pesticide or class, but these reductions would result in similar effects on ecosystem function overall (e.g., photosynthesis) (*10*). Second, communities respond consistently to pesticides within classes because of taxa-specific sensitivities to pesticides (*11*). Third, disruptions in ecosystem processes are mediated by changes to biodiversity.

We used 72 outdoor mesocosms to evaluate the effects of two control treatments (water and solvent), four simulated-pesticide treatments, and 12 pesticides on tri-trophic temperate pond communities. The pesticide treatments were nested in four classes (organophosphates, carbamates, chloroacetanilides, triazines) and two types (insecticides and herbicides) (Fig. 1a,b). To represent pesticide runoff following rainfall, pesticides were applied singly at the beginning of the experiment at standardized environmentally relevant concentrations calculated using U.S. EPA software. Simulated-pesticide treatments were top-down or bottom-up food web manipulations that attempted to mimic direct (i.e. lethal) effects of actual herbicides and insecticides on algae and zooplankton abundances, respectively. Mesocosm studies are an efficient approach to toxicity testing as they provide toxicity data on multiple species simultaneously under environmentally realistic conditions (Supplemental Text).

**Figure 1.**
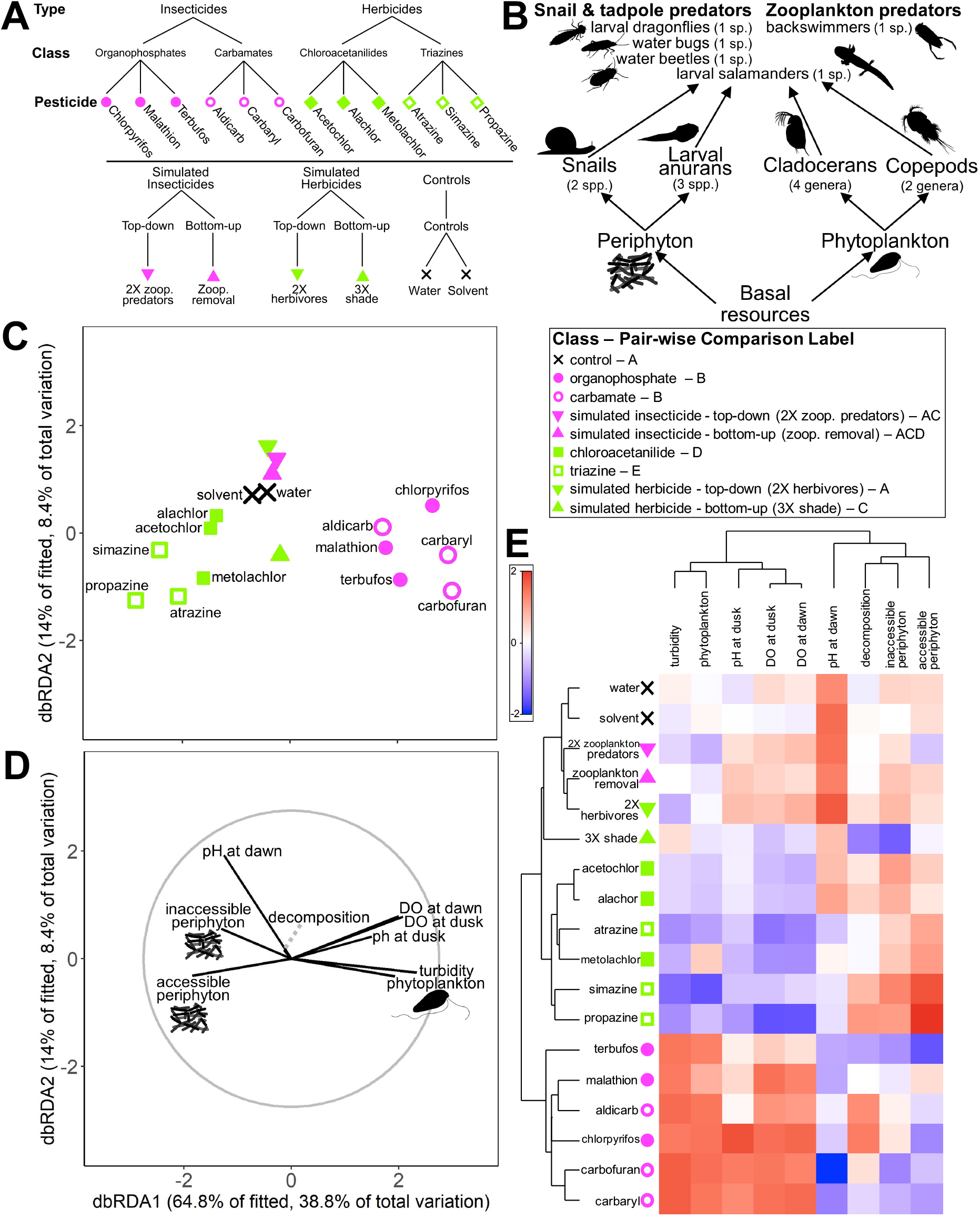
Experimental design and similarity of ecosystem responses by pesticide type. **A)** Food web diagram of experimental communities. **B)** Experimental design showing hierarchical structure of treatments. Each treatment was replicated four times with mesocosm as the replicate. **C)** Distance-based redundancy analysis (dbRDA) plot of multivariate ecosystem responses showing differences among treatments grouped by pesticide type. Individual points are the centroids of 18 treatments in the experiment. For pair-wise comparisons, treatments sharing letters are not different from each other. **D)** Vector overlay of ecosystem responses for the corresponding dbRDA plot. The gray circle corresponds to vector lengths that would have a correlation coefficient of one with each axis. **E)** Cluster diagram of experimental treatments and ecosystem-level responses showing grouping of experimental treatments according to pesticide type.

Pesticide type explained 46% of the variation in ecosystem function associated with the pesticide treatments (Table S1, Fig. 1c). Herbicides were associated with a decrease in suspended phytoplankton that led to increased abundance of attached periphyton, likely through an increase in light availability (Fig. 1d,e). With decreased phytoplankton, pH at dusk and dissolved oxygen at dawn and dusk decreased (Fig. 1d,e), reflecting reduced primary productivity relative to whole system respiration, which might have been driven by the decomposition of dead phytoplankton.

In contrast to herbicide-exposed systems, insecticide-exposed systems had an increase in phytoplankton, which shaded and thus decreased periphyton (Fig. 1d,e), an effect shown in other studies (*11*). Increases in phytoplankton were likely driven by shifts in zooplankton communities away from cladocerans and towards copepods (described below). The corresponding increase in dissolved oxygen at dawn and dusk and in acidity at dawn reflects increased respiration and primary production (Fig. 1d,e). While some variation in ecosystem responses was also explained by individual pesticides, it was small relative to variation explained by pesticide type and was driven by two pesticides (e.g., alidcarb within carbamates, chlorpyrifos within organophosphates; Fig. 1c).

We tested for the effects of individual pesticides, classes, and types separately on the single-trophic-level zooplankton community (six zooplankton genera) and on the tri-trophic community (insect and salamander predators, snail and anuran herbivores, and periphyton and phytoplankton primary producers). Similar to ecosystem function, pesticide type explained the majority of the variance (44.2%) in the zooplankton community, followed by pesticide class (18.8%) (Table S1). Distance-based redundancy analysis (dbRDA) showed that: 1) herbicide-and insecticide-treated mesocosms had distinct zooplankton communities, 2) within their respective pesticide types, organophosphate insecticides, chloroacetanilide herbicides, and triazine herbicides caused further distinction in zooplankton communities, and 3) there was relatively high multivariate dispersion within the carbamate class (Fig. 2a). In response to insecticides, cladoceran zooplankton were virtually eliminated, likely leading to competitive release of copepods (Fig. 2b,c). Reduced cladocerans, which are more efficient phytoplankton grazers than copepods (*12*), likely drove the increased relative abundance of phytoplankton in the algal community (Fig. 1c). In contrast to the changes in community composition associated with insecticides, herbicides decreased zooplankton abundance with no apparent change in composition (Fig. S3), likely by reducing phytoplankton (i.e., bottom-up effect). The stronger bottom-up effect of triazines compared to chloroacetanilide herbicides on zooplankton was likely because of longer environmental persistence (soil half-lives 110-146 d vs. 14-26 d, respectively [Pesticide Action Network Pesticide Database]). Thus, consistent with the ecosystem function results, these findings on the zooplankton community suggest that ecological risk assessment can be largely simplified to generalized effects of pesticide type or class.

**Figure 2.**
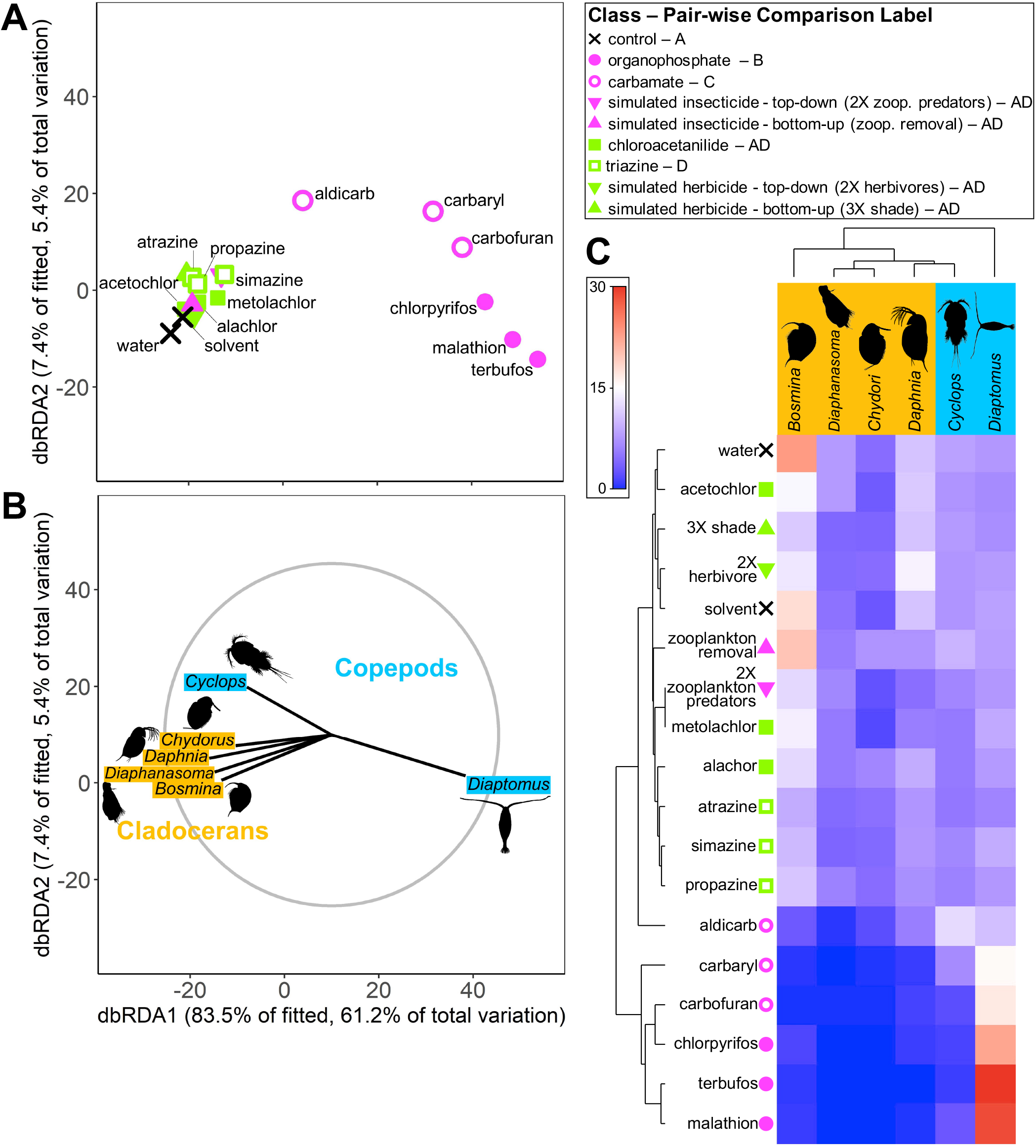
Zooplankton communities respond consistently to pesticides within type, class, and individual pesticide. **A)** dbRDA plot, **B)** vector overlay, and **C)** cluster diagram of multivariate zooplankton densities showing differences among type, class, and individual pesticide.

In the tri-trophic community, variation explained by pesticides was about equally distributed among type, class, and individual pesticide (Table S1). The dbRDA showed that: 1) herbicide-and insecticide-treated mesocosms had distinct communities, 2) within herbicides, triazines and chloroacetanilides classes caused further distinction in communities, and 3) there was relatively high multivariate dispersion in communities exposed to carbamate and organophosphate insecticides (Fig. 3a). Overall, survival of predators was low with insecticides, except for aldicarb (Fig. 3b,c). Amphibian and snail prey generally had greater positive responses to insecticides compared to controls or herbicides (Fig. 3b,c), suggesting benefits from a release from predators, a trend found in other studies (*13*). When we grouped responses of taxa by functional role in the community (algae, herbivores, and predators), the amount of variance accounted for by pesticide type (29%) was almost double the variance accounted for by either pesticide class (17.6%) or individual pesticide (17.3%) (Table S1). From these results we conclude that complexity in predicting the effects of pesticides on communities could be reduced by evaluating responses of functional groups instead of individual taxonomic responses.

**Figure 3.**
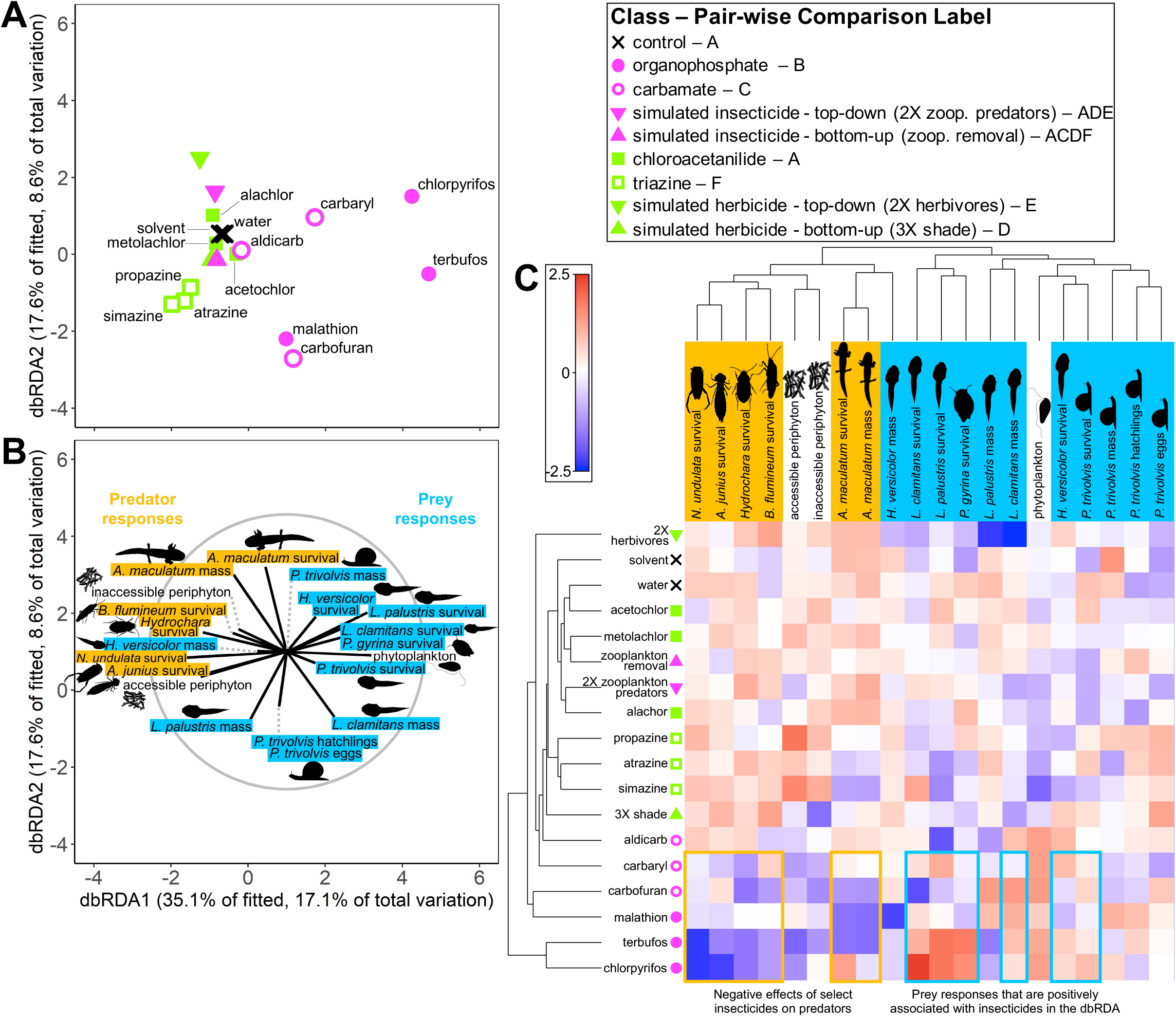
Insecticides generally reduce insect predators in tri-trophic communities, resulting in increases in the survival and growth of their prey. **A)** dbRDA plot, **B)** vector overlay, and **C)** cluster diagram of multivariate community responses showing differences among type, class, and individual pesticide. When responses within tri-trophic communities are grouped by functional role (algae, herbivores, predators), type explains twice as much variation as class and individual pesticide (Table S1).

Next, we evaluated if the effects of pesticides occurred via changes in abundance of ‘targeted taxa’ by examining simulated pesticide treatments. This evaluation was motivated by a call to uncover non-target effects and mechanisms by which anthropogenic drivers alter biodiversity and ecosystem function via direct manipulations of taxa (*6*). With the exception of the bottom-up simulated herbicide, the effects of simulated pesticide treatments on ‘targeted taxa’ did not match the effects of pesticide classes (pair-wise comparisons Fig. 1–3), likely because of difficulties in sustaining manipulations that matched the magnitude and specificity of actual pesticides (see Supplemental Text and Fig. S3 for details) on species with short generation times. While manipulating species composition has been critical historically in the study of biodiversity-ecosystem function, the same approaches are likely not well-suited for 1) species that can exhibit population dynamics in the timescale of an experiment because they should be able to quickly recover from a manipulation and 2) studies of disturbance because matching the complexity of the effects of actual disturbance is challenging.

Only the bottom-up simulated herbicide (i.e., triple shade cloth) was effective at reducing the targeted taxa, as it significantly reduced algae (t=2.009, *p*=0.015, Fig. 1). When we compared this treatment to triazine herbicides, there were minor differences in effects on ecosystem function and community composition, but only in magnitude, not direction, and neither triazines nor bottom-up simulated herbicides led to detectable lethal effects on non-target taxa. Bottom-up simulated herbicides and triazines grouped together in the tri-trophic community cluster diagram (Fig. 3c), and triazines were not different from bottom-up simulated herbicide treatments in the zooplankton community analysis (Fig. 2c). These results suggest that the observed effects of triazines on freshwater ecosystems were driven predominantly by direct toxicity to targeted taxa and their associated indirect effects, rather than non-target effects.

Path models revealed that effects of herbicides on ecosystem functions, such as primary productivity and respiration, were mediated by their effects on biodiversity (Fig. 4a; via changes in evenness of algae and not richness, Fig. S5). In contrast, although insecticides influenced biodiversity (Fig. 4c), insecticides primarily altered ecosystem function via the direct path, which captures both direct effects of pesticides (e.g., biogeochemical alterations to carbon cycle) and indirect effects mediated by unmeasured aspects of biodiversity (e.g., microbiota). The absence of a significant biodiversity-mediated effect of insecticides is likely because insecticides directly reduced species (e.g., predators, zooplankton) that contributed little to the ecosystem functions we measured (photosynthesis, respiration, decomposition). Given that species responses showed greater variability to pesticides compared to ecosystem functions and pesticide-induced changes to ecosystem functions were at times mediated by changes to biodiversity, the observed consistency in the responses of ecosystem functions to pesticides within pesticide types is likely driven by functional redundancies of species. Indeed, when we simplified our community by functional roles, pesticide type explained almost twice the variance as pesticide class or individual pesticide (Table S1).

**Figure 4.**
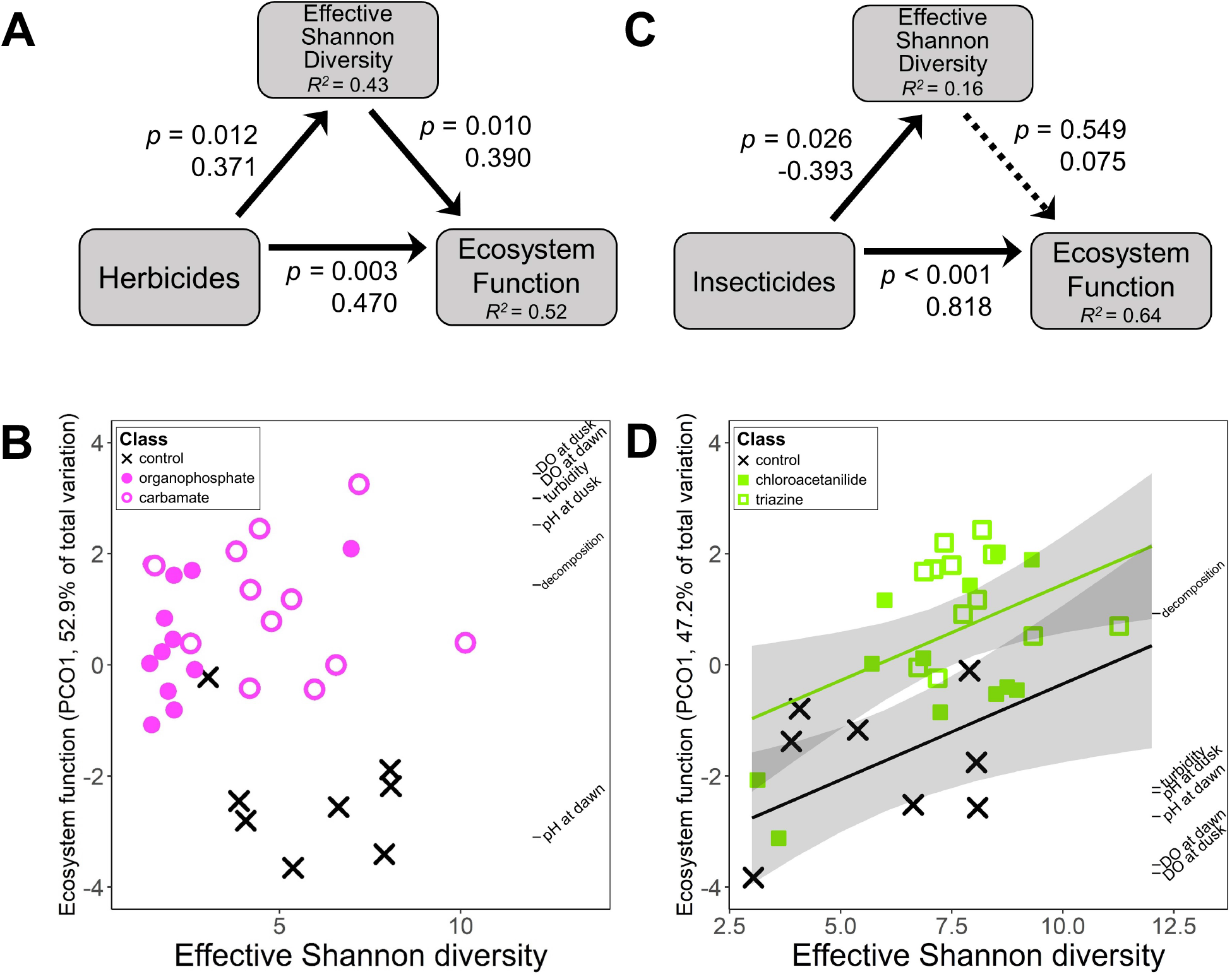
Path models among pesticides, biodiversity, and ecosystem function. Relationships among Shannon diversity, ecosystem functions, and **A)** herbicides or **C)** insecticides, and associated scatterplots of Shannon diversity and ecosystem functions for the **B)** herbicides and controls and **D)** insecticides and controls. Herbicides affected ecosystem function by increasing evenness and reducing primary productivity, whereas there was no detectable relationship for insecticides. Solid arrows are significant paths, and dotted arrows are non-significant paths. P-values, standardized coefficients, and conditional R^2^ values are provided.

Ecological risk assessment, made complex by tens of thousands of synthetic chemicals and diverse species assemblages, could be simplified by assessing groups of chemicals that share similar chemical structures or environmental targets and examining responses of functional groups of taxa (*14*). A similar approach is taken by Quantitative Structure-Activity Relationships (QSAR) (*15*), which attempts to assess toxicity of synthetic chemicals using information on compound structure. The clustering of pesticides in this study was not consistent with predictions from QSAR models, likely because predictions are based on few model organisms and do not include indirect effects (Supplemental Text). Nevertheless, this study suggests that simplifying groups of commercial chemicals that share similar chemical structures or environmental targets can be extended from individuals to ecosystems, which would improve predictions and provide more time and resources to identify potential exceptions to general patterns. Before these changes are adopted by risk managers, additional testing would be warranted to evaluate if the patterns observed here are consistent across additional contaminants, biomes, and community compositions and structures.

## Funding

Funding was provided by the National Institutes of Health (R01GM109499, R01TW010286-01), the National Science Foundation (EEID-1518681, EF-1241889, IOS-1754868, IOS-1651888), and the Department of Agriculture (NRI 2009-35102-0543).

## Author contributions

J.T.H, T.R.R, and J.R.R. designed the experiment, J.T.H, T.R.R., and H.J.C, conducted the experiment, S.L.R., M.B.M, and J.R.R. conducted the analyses, S.L.R. wrote the manuscript, and all authors contributed to editing the manuscript.

## Competing interests

The authors declare no competing interests.

## Data and materials availability

All data are available at Figshare under the **accession number** XXXXXXX.

## References

1. J. S. Baron, N. L. Poff, P. L. Angermeier, C. N. Dahm, P. H. Gleick, N. G. Hairston, R. B. Jackson, C. A. Johnston, B. D. Richter, A. D. Steinman, Meeting ecological and societal needs for freshwater. Ecol. Appl. 12, 1247–1260 (2002).

2. E. S. Bernhardt, E. J. Rosi, M. O. Gessner, Synthetic chemicals as agents of global change. Front. Ecol. Environ. 15, 84–90 (2017).

3. N. T. Baker, W. W. Stone, “Estimated annual agricultural pesticide use for counties of the conterminous United States, 2008-12: U.S. Geological Survey Data Series 907” (U.S. Geological Survey, Reston, VA), p. 9.

4. S. L. Rumschlag, N. T. Halstead, J. T. Hoverman, T. R. Raffel, H. J. Carrick, P. J. Hudson, J. R. Rohr, Effects of pesticides on exposure and susceptibility to parasites can be generalised to pesticide class and type in aquatic communities. Ecol. Lett. 22, 962–972 (2019).

5. T. A. McMahon, N. T. Halstead, S. Johnson, T. R. Raffel, J. M. Romansic, P. W. Crumrine, J. R. Rohr, Fungicide-induced declines of freshwater biodiversity modify ecosystem functions and services. Ecol. Lett. 15, 714–722 (2012).

6. F. De Laender, J. R. Rohr, R. Ashauer, D. J. Baird, U. Berger, N. Eisenhauer, V. Grimm, U. Hommen, L. Maltby, C. J. Meliàn, F. Pomati, I. Roessink, V. Radchuk, P. J. Van den Brink, Reintroducing environmental change drivers in biodiversity–ecosystem functioning research. Trends Ecol. Evol. 31, 905–915 (2016).

7. B. J. Cardinale, D. S. Srivastava, J. E. Duffy, J. P. Wright, A. L. Downing, M. Sankaran, C. Jouseau, Effects of biodiversity on the functioning of trophic groups and ecosystems. Nature. 443, 989 (2006).

8. K. N. Suding, S. Lavorel, F. S. Chapin, J. H. C. Cornelissen, S. Díaz, E. Garnier, D. Goldberg, D. U. Hooper, S. T. Jackson, M.-L. Navas, Scaling environmental change through the community-level: a trait-based response-and-effect framework for plants. Glob. Change Biol. 14, 1125–1140 (2008).

9. D. Tilman, F. Isbell, J. M. Cowles, Biodiversity and ecosystem functioning. Annu. Rev. Ecol. Evol. Syst. 45, 471–493 (2014).

10. J. W. Fleeger, K. R. Carman, R. M. Nisbet, Indirect effects of contaminants in aquatic ecosystems. Sci. Total Environ. 317, 207–233 (2003).

11. J. Hua, R. Relyea, Chemical cocktails in aquatic systems: pesticide effects on the response and recovery of >20 animal taxa. Environ. Pollut. Barking Essex 1987. 189, 18–26 (2014).

12. U. Sommer, F. Sommer, Cladocerans versus copepods: the cause of contrasting top-down controls on freshwater and marine phytoplankton. Oecologia. 147, 183–194 (2006).

13. S. D. Peacor, E. E. Werner, Predator Effects on an Assemblage of Consumers Through Induced Changes in Consumer Foraging Behavior. Ecology. 81, 1998–2010 (2000).

14. J. R. Rohr, C. J. Salice, R. M. Nisbet, The pros and cons of ecological risk assessment based on data from different levels of biological organization. Crit. Rev. Toxicol. 46, 756–784 (2016).

15. P. Mazzatorta, E. Benfenati, P. Lorenzini, M. Vighi, QSAR in ecotoxicity: an overview of modern classification techniques. J. Chem. Inf. Comput. Sci. 44, 105–112 (2004).

16. J. Travis, H. M. Wilbur, in Ecological communities: conceptual issues and evidence (Princeton, New Jersey, USA, 1984), pp. 113–122.

17. J. R. Rohr, A. M. Schotthoefer, T. R. Raffel, H. J. Carrick, N. Halstead, J. T. Hoverman, C. M. Johnson, L. B. Johnson, C. Lieske, M. D. Piwoni, P. K. Schoff, V. R. Beasley, Agrochemicals increase trematode infections in a declining amphibian species. Nature. 455, 1235–1239 (2008).

18. L. Jost, Entropy and diversity. Oikos. 113, 363–375 (2006).

