## Supplement for "Consistent Effects of Pesticides on Community Structure and Ecosystem Function in Freshwater Systems"

**This PDF file includes:**

Materials and Methods
Supplemental Text
Figs. S1 to S5
Tables S1and S2
References (*16-18*)

### Materials and Methods

Experimental Design and Community Composition

We conducted a randomized-block experiment at the Russell E. Larsen Agricultural Research Center (Pennsylvania Furnace, PA, USA) with replicated mesocosm ponds. Mesocosms were 1,100-L cattle tanks covered with 60% shade cloth. The spatial block was distance from a tree line in our mesocosm field. Three weeks before pesticide application, these mesocosms were filled with 800 L water, 300 g mixed hardwood leaves, and inoculations of zooplankton, periphyton, and phytoplankton homogenized from four local ponds. Just before pesticide application on the same day, each tank received two snail, three larval anuran, one larval dragonfly, one water bug, one water beetle, one larval salamander, and one backswimmer species (11 *Helisoma* (*Planorbella*) *trivolvis*, 10 *Physa gyrina*; 20 *Hyla versicolor*, 20 *Lithobates palustris*, 20 *Lithobates clamitans*; 2 *Anax junius*; 2 *Belostoma flumineum*; 5 *Hydrochara* sp.; 3 *Ambystoma maculatum*; 6 *Nototeca undulata*) (Fig. 1a). These community members naturally coexist and were applied at naturally occurring densities (*16*). Initial conditions of some mesocosms varied in simulated pesticide treatments (see below).

We randomly assigned 18 treatments (12 pesticides, 4 simulated pesticides, 2 controls) with four replicate mesocosms of each treatment, which resulted in 72 total mesocosms (Fig. S1b). The 12 pesticide treatments were nested; we included two pesticide types (insecticide, herbicide), two classes within each pesticide type (organophosphate insecticide, carbamate insecticide, chloroacetanilide herbicide, triazine herbicide), and three different pesticides in each of four classes (Fig. S1b). To represent runoff of pesticides into freshwater systems following a rainfall event, we applied single doses of technical grade pesticides at environmentally relevant concentrations at the beginning of the experiment. To ensure our exposures represented environmental relevance, we used estimated environmental concentrations of pesticides, calculated by U.S. Environmental Protection Agency's GENEEC v2 software, Table S2). Our design also included water and solvent (0.0001% acetone) controls (Fig. 1b). Pesticides were obtained from ChemService (West Chester, PA, USA). Nominal concentrations of pesticides (μg/L) were: 64 chlorpyrifos, 101 malathion, 171 terbufos, 91 aldicarb, 219 carbaryl, 209 carbofuran, 123 acetochlor, 127 alachlor, 105 metolachlor, 102 atrazine, 202 simazine, and 106 propazine. We collected composite water samples one hour after application to mesocosms and shipped samples on ice to Mississippi State Chemical Laboratory to verify these nominal concentrations. Measured concentrations of pesticides (μg/L) were: 60 chlorpyrifos, 105 malathion, 174 terbufos, 84 aldicarb, 203 carbaryl, 227 carbofuran, 139 acetochlor, 113 alachlor, 114 metolachlor, 117 atrazine, 180 simazine, and 129 propazine.

The four simulated pesticide treatments were top-down or bottom-up food web manipulations intended to mimic effects of actual herbicides and insecticides on community members. These manipulations occurred once and were concurrent with the timing of pesticide applications. Top-down and bottom-up simulated insecticide treatments were designed to reduce densities of zooplankton, simulating effects of insecticides on zooplankton survival. For top-down simulated insecticides, we doubled the densities of zooplankton predators by including six total *A. maculatum* larval salamanders and 12 *N. undulata* backswimmers per mesocosm. For bottom-up simulated insecticides (i.e., direct manipulation of a lower arthropod trophic level), we removed zooplankton with a vertical tow-net. Top-down and bottom-up simulated herbicides were designed to reduce algae, simulating effects of herbicides on survival and growth of algae. For top-down simulated herbicides, we doubled the densities of large herbivores to increase grazing pressure by including 22 *H. trivolvis* snails, 20 *P. gyrina* snails, 40 *H. versicolor* larval anurans, 40 *L. palustris* larval anurans, and 40 *L. clamitans* larval anurans per mesocosm. For bottom-up simulated herbicides, we covered mesocosms in three sheets of 60% shade cloth in an attempt to block light and reduce photosynthesis. The experiment ran for four weeks, from June to July.

Measurements of Experimental Responses

During the experiment, we sampled periphyton using clay tiles (100 $cm^2$) oriented perpendicularly along the bottom of the mesocosm. Each mesocosm had two periphyton measurements: 'inaccessible periphyton' taken from caged clay tiles that excluded herbivores and 'accessible periphyton' taken from clay tiles that were uncaged allowing herbivore access. We sampled phytoplankton from water samples taken 10 cm below the water surface. Periphyton was scrubbed from tiles and phytoplankton from water samples (10 mL) were filtered onto glass fiber filters (under low vacuum pressure, <10 psi; Whatman EPM 2000, 0.3 μm, 47 mm) to estimate associated chlorophyll concentrations. The chlorophyll concentration of each filter was determined using an organic extraction procedure with a 50:50 mixture of 90% acetone to DMSO. We measured chlorophyll-*a* concentrations using a standard fluorometric technique. We scored water clarity, a metric of light availability, on a scale from one (clear) to five (opaque) blind to treatment. We measured pH and dissolved oxygen (DO) at dusk and dawn on subsequent days using hand-held meters (YSI, Yellow Springs, OH, USA). We measured decomposition by taking the dry mass of hardwood leaf packets in each mesocosm at the beginning and the end of experiment. In addition, we sampled snail egg masses and hatchlings using two rectangular pieces of Plexiglass (465 $cm^2$) in each mesocosm, one hung on the side and one on the bottom of the mesocosm. Zooplankton were collected from the entire water column by placing a PVC pipe (10 cm diameter, 60 cm height) upright in the center of each tank, capping the bottom, and pouring the water through a 20 μm Nitex mesh. We collected two samples of zooplankton from each mesocosm, and we combined and preserved the samples in 70% ethanol. Zooplankton were counted and identified in 5 mL subsamples for each mesocosm using a zooplankton counting wheel (Wildlife Supply Company, Yulee, FL, USA) and a dissecting microscope. At the end of the experiment, mesocosms were drained, and the remaining animals were counted, euthanized, and preserved. Two previous manuscripts, which use the same design as the current manuscript, also describe this experimental design and methods (*4*, *17*).

Statistical Analyses

To test for the consistency of effects of type, class, and individual pesticide on aquatic ecosystem processes and communities and to attribute the variation explained to each pesticide level of organization while accounting for the nested structure of our experimental design (Figure 1b), we completed permutational analyses of variance (PERMANOVA). For nested PERMANOVA models, the predictors were the following random categorical terms: type (insecticide, herbicide), class (carbamate, organophosphate, chloroacetanilide, triazine) nested within type, and pesticide (12 in total) nested within class within type. These models did not include controls or simulated pesticides because these treatments were not hierarchically nested (Fig. S1b). We evaluated 9999 permutations using residuals under a reduced model. Following nested PERMANOVAs, we used pair-wise multiple comparisons tests using PERMANOVAs to evaluate differences among controls, organophosphates, carbamates, top-down simulated insecticides, bottom-up simulated insecticides, chloroacetanilides, triazines, top-down simulated herbicides, and bottom-up simulated herbicides. In these pair-wise comparisons, we evaluated

9999 unrestricted permutations of raw data. All PERMANOVAs also included spatial block as a random predictor to account for variation in sunlight associated with distance from a tree line. Preliminary analyses showed that exclusion of the block did not change the results. In all PERMANOVAs, test statistics associated with Type III partial sums of squares were evaluated.

We conducted four nested PERMANOVAs. Our first nested PERMANOVA focused on ecosystem processes and included the following responses: pH and dissolved oxygen taken both at dawn and dusk, decomposition (percent mass loss of hardwood leaf packets), turbidity (water clarity scores from 1 to 5), and densities of phytoplankton, accessible periphyton, and inaccessible periphyton (measured via chlorophyll-*a*). The resemblance matrix for these responses was constructed using a Euclidean distance matrix of log-transformed and normalized values.

Our second and third nested PERMANOVAs focused on community structure. We separated community members into two statistical models based on the forms of response variables; those whose response variables were densities based on counts (zooplankton) and those community members whose response variables were survival, mass, reproductive rates, or density abstracted from chlorophyll measurements (insect predators, snail and tadpole herbivores, and algae; termed the tri-trophic community). The multivariate response for the zooplankton community included densities of *Daphnia*, *Diaphanasoma*, *Chydorus*, *Bosmina*, *Diaptomus*, and *Cyclops*. Zooplankton community analyses were based on square-root transformed densities using Bray-Curtis similarities. The multivariate response for the tri-trophic community model included: survival (0 to 1) of all amphibian, snail, and insect community members; average mass of surviving individuals for each amphibian species and *H. trivolvis* snails; average number of hatchlings and eggs per surviving *H. trivolvis* snail; and densities of phytoplankton and periphyton. Mass and reproductive rates were standardized to the number of surviving individuals to account for the different densities added to each tank at the beginning of the study (i.e. extra herbivores in top-down simulated herbicide treatment and extra predators in bottom-up simulated insecticide treatment). Mass and reproductive rates of *P. gyrina* were not included because of low survival across treatments. Survival rates were arc-sine square-root transformed and normalized, and all other variables were log-transformed and normalized. Tri-trophic community analyses were based on Euclidean distances.

Finally, our fourth nested PERMANOVA evaluated a simplified tri-trophic community. We simplified the tri-trophic community responses into three functional roles within the community: algae, herbivores, and predators. Tri-trophic community responses of individua taxa were transformed and normalized as described previously, and then they were averaged according to functional group. We averaged densities of periphyton and phytoplankton into a single "algae" response, all amphibian and snail responses into a single "herbivore" response, and all insect and salamander responses into a single "predator" response. The simplified tri-trophic community model was based on Euclidean distances.

To visualize consistency of effects within type, class, and individual pesticides on multivariate ecosystem and community responses and to compare pesticide effects to simulated pesticides and controls, we used distance-based redundancy analyses (dbRDA) and two-way cluster diagrams. The dbRDAs were based on appropriate resemblance matrices for ecosystem and community responses as described above. The underlying categorical predictors in the models for ecosystem processes and tri-trophic communities included: the spatial block, organophosphate, carbamate, chloroacetanilide, triazine, top-down simulated insecticide, bottom-up simulated insecticide, top-down simulated herbicide, bottom-up simulated herbicide, and control. In the zooplankton analyses, all previous predictors were included except for spatial block because it was not significant in the PERMANOVA test. In the dbRDA plots, when spatial block was included in the ecosystem and tri-trophic community plots, we show the centroid values for the 18 experimental replicates.

As an alternative to the dbRDAs presented in the main text, we also visualized the consistency of effects within type, class, and individual pesticide on ecosystem, tri-trophic community, and zooplankton responses and compared pesticide effects to simulated pesticides and controls, using principal coordinates analyses (PCoA) (Figs. S2, S3, S5). PCoAs were based on appropriate resemblance matrices as described previously. PCoAs were conducted in PERMANOVA+ for PRIMER and resulting data were exported. Point and vector plots were made using exported data and the '*ggplot2*' package in R. Ellipses on point plots represent 95% confidence intervals of groups based on standard errors and were made using the ordiellipse function in the '*vegan*' package.

For the two-way cluster diagrams, clusters of pesticide treatments were based on centroid distances of the appropriate resemblance matrices. Clusters of multivariate responses were based on Euclidean distance resemblance matrices of averaged treatment responses. Before averaging, ecosystem and community responses were transformed and normalized as described previously. In clustering of treatments and responses, the cluster mode was the group average. In the PERMANOVAs for ecosystem processes and tri-trophic community responses, the effect of block was significant (Table S1). Thus, we accounted for the effect of block by taking the residuals of simple linear regressions with individual ecosystem or tri-trophic community responses as the independent variable and block as the predictor in the generation of the shaded values of the two-way cluster diagrams. Then, we averaged these block-adjusted treatment responses with the 'shade plot' function in PRIMER. For the zooplankton community, the effect of block was not significant in the PERMANOVA model (Table S1), so shaded values of the two-way cluster diagrams were simply the averaged treatment responses. All PERMANOVA models, pair-wise comparisons, dbRDAs, and two-way cluster diagrams were executed using PERMANOVA+ for PRIMER version 7 (PRIMER-E Ltd, Plymouth, UK). For ease of visualization of dbRDA and PCoA plots, data from PERMANOVA+ for PRIMER were exported, and plots were made using '*ggplot2*' package in R.

To compare the level of support for direct versus indirect biodiversity-mediated effects by which pesticides might influence ecosystem processes, we performed path analyses using the '*piecewiseSEM'* package. We chose to use path analyses because they allowed us to test multiple linked hypotheses via the consideration of multiple variance-covariance matrices in which variables serve as both dependent and independent variables. In evaluating the effects of herbicides and insecticides on ecosystem processes, we chose to compare two mechanistic paths: the direct effects of herbicides or insecticides on ecosystem processes and the indirect effects in which the effects of herbicides or insecticides are mediated by the impact of biodiversity on ecosystem processes. The unit of replication was the mesocosm, and each path model contained 32 independent replicates. Within the path models, we accounted for the effect of spatial block by using linear mixed effect models in which block was the random intercept term. For our biodiversity metrics, we calculated Hill numbers of species richness ($q = 0$), Shannon diversity ($q = 1$, exponent of Shannon index), and Simpson diversity ($q = 2$, inverse of Simpson index) (*18*). Hill numbers are preferred over other diversity metrics because units of Hill numbers are effective number of species as opposed to unitless metrics that are challenging to interpret (*18*). Biodiversity metrics were calculated in PRIMER. We present the results of path analyses using

Shannon diversity in the main text, while results using species richness and Simpson diversity are included in the Supplemental Information (Fig. S6). For all path models, ecosystem function is the first axis from a principal coordinates analysis of the Euclidean resemblance matrix of log-transformed and normalized ecosystem responses including: pH and dissolved oxygen taken both at dawn and dusk, decomposition, and turbidity. Fit statistics indicate that all path models fit well (Fisher's C = 0, $p$-value = 1).

### Supplemental Text

#### Costs and Benefits of Mesocosm Studies

Using mesocosms studies in toxicity testing has previously been criticized because of high costs compared to traditional single species toxicity tests like the LC50. While a mesocosm study takes more time and money to conduct compared to a single LC50 study, it also provides information on the toxicity to multiple organisms under more environmentally realistic conditions. To properly consider the costs and benefits of a mesocosm experiment, an estimate would need to consider both the abundance of toxicity data and the ecological realism of the data. For instance, assume that the average mesocosm study provides toxicity information for 10 species, then this study should be compared to the costs of conducting 10 LC50 studies. Additionally, the toxicity data from the mesocosm experiment should be of higher value because it includes realistic ecological complexities (e.g. direct and indirect effects, recovery dynamics of the populations and communities). In comparison, the LC50 study measures the toxicity of a single organism under contrived lab conditions. When these two approaches are more appropriately compared, the benefits of mesocosm experiments could outweigh the costs in comparison to traditional toxicology studies.

#### Performance of Simulated Pesticide Treatments

With the exception of bottom-up simulated herbicide, the effects of simulated pesticides did not match the effects of pesticide classes (pair-wise comparisons Fig. 1-3). These treatments performed poorly likely because manipulating taxa did not match the magnitude or the specificity of the long-term effect of pesticides. Top-down simulated herbicides (i.e., doubled herbivores) were designed to reduce algae, but the added tadpoles and snails mostly feed on periphyton, while the actual herbicides had a greater long-term net negative effect on phytoplankton (Fig. 1). Top-down (i.e., doubled zooplankton predators) and bottom-up (i.e., zooplankton removal) simulated insecticides both failed to replicate the differential toxicity that insecticides had on cladoceran versus copepod zooplankton (Fig. S3).

#### Evaluation of Acute Aquatic Toxicity Using QSAR Approaches

We completed analyses to evaluate if the consistency of pesticides on aquatic systems observed in the current study could be predicted by QSAR methods based on the structure of the pesticides alone. We used the QSAR Toolbox (https://qsartoolbox.org/) developed in partnership with The Organisation for Economic Co-operation and Development (OECD). QSAR Toolbox is a centralized, open-source software system for predicting toxicity of chemicals by applying a category approach. One functionality of the software is the clustering of chemicals into similar groups based on the predicted acute aquatic toxicity. The predicted acute aquatic toxicity is based solely on the pesticide's chemical structure. We conducted three different analyses that examined the clustering of our 12 pesticides. Below, we describe the three different analyses and the results.

1) Acute Aquatic Toxicity Classification by Verhaar
Description: The Acute aquatic toxicity classification by Verhaar consists of parametric and structural rules to mimic the Verhaar rules developed by Toxtree software. This system is introduced for chemical categorization purposes or can be used for the prioritization of chemicals for subsequent testing.

Results:
Group 1: All carbamates and organophosphates
Group 2: All triazines
Group 3: All chloroacetanilides

2) Acute Aquatic Toxicity by Mode of Action by OASIS
Description: This profile divides chemicals in different categories according to their acute toxic mode of action (MOA). 2D structural information is used only to identify the MOA of chemicals. Based on theoretical and empiric knowledge the following seven hierarchically ordered MOA are distinguished: Aldehydes; alpha, beta-Unsaturated alcohols; Phenols and Anilines; Esters; Narcotic Amines; Basesurface narcotics.

Results:
Group 1: All carbamates, organophosphates, and chloroacetanilides
Group 2: All triazines

3) Aquatic Toxicity Classification by ECOSAR
Description: The Aquatic Toxicity Classification by ECOSAR profiler consists of molecular definitions to mimic the structural definitions of chemical classes within the U.S. Environmental Protection Agency's Ecological Structure-Activity Program (ECOSAR™). ECOSAR™ contains a library of class-based structural activity relationships for predicting aquatic toxicity, overlaid with an expert decision tree based on expert rules for selecting the appropriate chemical class for evaluation of the compound.

Results:
Group 1: All triazines
Group 2: All chloroacetanilides
Group 3: Carbofuran and carbaryl (both carbamates)
Group 4: Aldicarb (carbamate)
Group 5: Terbufos (organophosphate)
Group 6: Malathion (organophosphate)
Group 7: Chlorpyrifos (organophosphate)

So, the result is that the pesticide groups vary based on the underlying assumptions of the model even though all three models are generally trying to predict pesticides that have similarity in the aquatic toxicities. These groupings are not very consistent with the observed toxicities to taxa in our study (tri-trophic and zooplankton communities). For instance, in the tri-trophic community analyses, the pairwise comparisons would suggest that all four pesticide classes behave differently. In the analyses of the zooplankton, the pairwise comparisons suggest that chloroacetanilides and triazines should group together while organophosphates and carbamates should form two additional separate groups.

The main issues with these groups of pesticides based on predicted toxicity is that they do not consider the variation across responses of groups of taxa and they do not consider indirect effects of pesticides. For instance, a QSAR model might predict direct effects of herbicides on algae, but that model will not include the indirect effect of herbicides on total zooplankton abundance.

Attribution of Silhouette Images

Silhouettes of organisms used throughout the manuscript are presented in accordance with licensing agreements. Below, we provide information on the creators' contributions to the images. Licensing agreements include: Public Domain Dedication 1.0 (https://creativecommons.org/publicdomain/zero/1.0/), Creative Commons Attribution 3.0 Unported license (https://creativecommons.org/licenses/by/3.0/), Attribution-ShareAlike 3.0 United States (https://creativecommons.org/licenses/by-sa/3.0/us/), and Attribution-ShareAlike 2.0 Generic (https://creativecommons.org/licenses/by-sa/2.0/).

Periphyton: created by Matt Crook, CC BY 3.0, no changes were made
Phytoplankton: created by T. Michael Keesey, CC0 1.0
*Helisoma* (*Planorbella*) *trivolvis*: created by Scott Hartman, CC0 1.0
*Physa gyrina*: created by Michael Mahon (vectorization), N. Yotarou (photography), CC BY 3.0
Anuran tadpole: created by Michael Mahon (vectorization), J.J. Harrison (photography), CC BY-SA 3.0 US
*Bosmina*: created by Michael Mahon (vectorization), S.F. Harmer and A.E. Shipley (photography), CC0 1.0
*Diaphanosoma*: created by Michael Mahon (vectorization), Museo Civico di Storia Naturale di Genova (photography), CC0 1.0
*Chydori*: created by Michael Mahon (vectorization), Henry Baldwin Ward and George Wipple (photography), CC0 1.0
*Daphnia*: not credited, CC0 1.0
*Cyclops*: created by Michael Mahon (vectorization), Great Lakes Image Collection (photography), CC0 1.0
*Diaptomus*: created by Michael Mahon (vectorization), NOAA Great Lakes Environmental Research Lab (photography), CC BY-SA 2.0
*Anax junius*: created by Michael Mahon (vectorization), J.J. Harrison (photography), CC BY 3.0
*Belostoma flumineum*: created by Dave Angelini, CC BY 3.0
*Hydrochara*: created by T. Michael Keesey (vectorization), Yves Bousquet (photography), CC BY 3.0
*Ambystoma maculatum*: created by Jake Warner, CC0 1.0
*Nototeca undulata*: created by Michael Mahon (vectorization); Christopher Johnson (photography), CC0 1.0

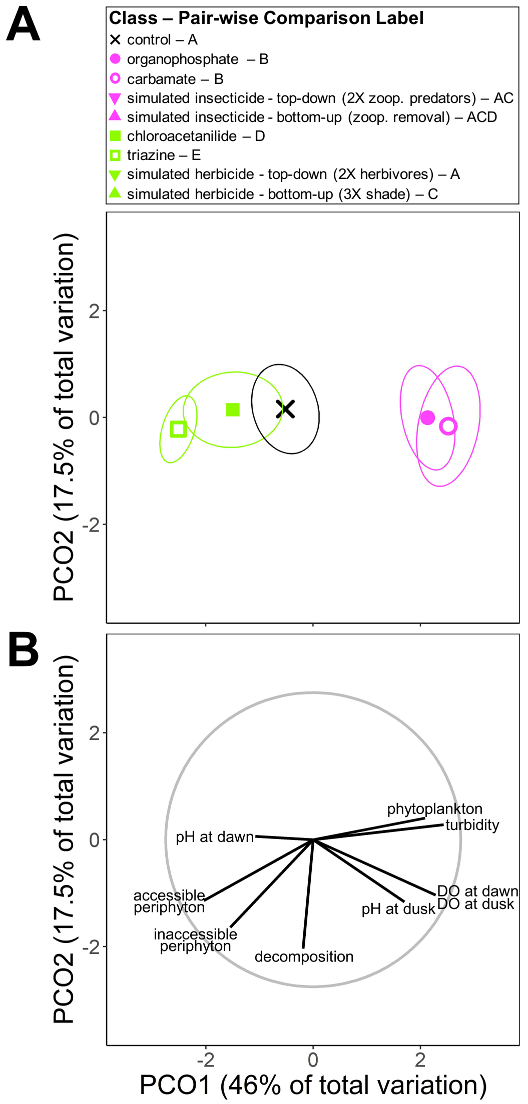


**Fig. S1 Principal coordinates analysis for multivariate ecosystem responses.** **A)** Principal coordinates analysis plot of multivariate ecosystem-level responses showing differences in pesticide treatments by type. Individual points are centroids for the representative treatment with ellipses based on 95% confidence intervals calculated using a standard error. All simulated herbicides and insecticides were different from corresponding pesticide treatments, so they were not included in the plot for ease of viewing. Pair-wise comparison labels are given in the figure legend. Treatments sharing letters are not different from each other. **B)** Vector overlay of log-transformed and normalized ecosystem-level responses for corresponding principal coordinates analysis plot. Gray circle shows relative vector distance lengths; the gray circle corresponds to vector lengths that would have a correlation coefficient of one.

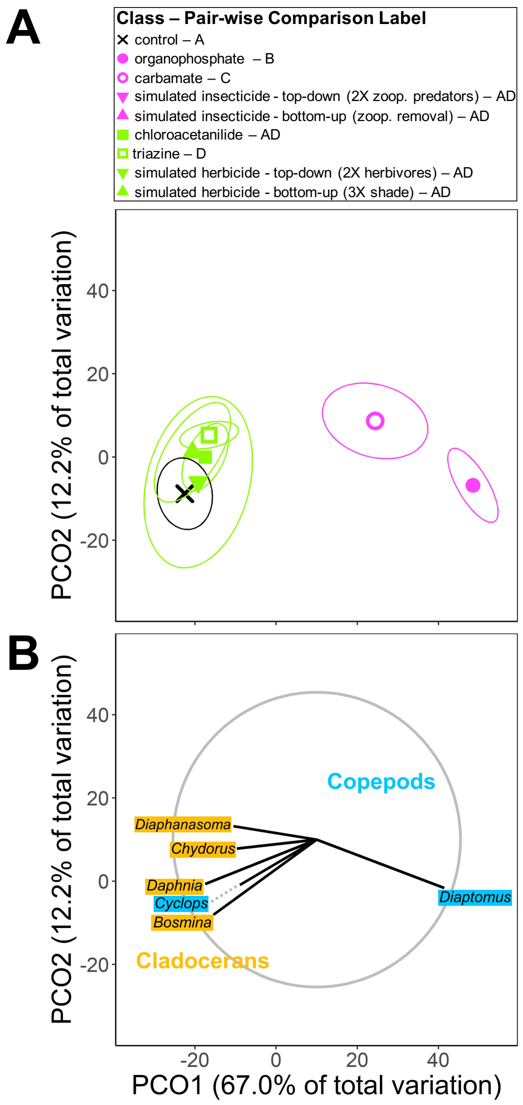


**Fig. S2 Principal coordinates analysis for multivariate zooplankton responses.** **A)** Principal coordinates analysis plot of multivariate zooplankton densities by genera showing differences in pesticide treatments by type, class, and individual pesticide. Individual points are centroids for the representative treatment with ellipses based on 95% confidence intervals calculated using a standard error. Simulated pesticides that were different from corresponding pesticide treatments, including bottom-up and top-down simulated insecticides, were not included in the plot for ease of viewing. Pair-wise comparison labels are given in the figure legend. Treatments sharing letters are not different from each other. **B)** Vector overlay of square-root transformed zooplankton densities by genera for corresponding principal coordinates analysis plot. Gray circle shows relative vector distance lengths; the gray circle corresponds to vector lengths that would have a correlation coefficient of one.

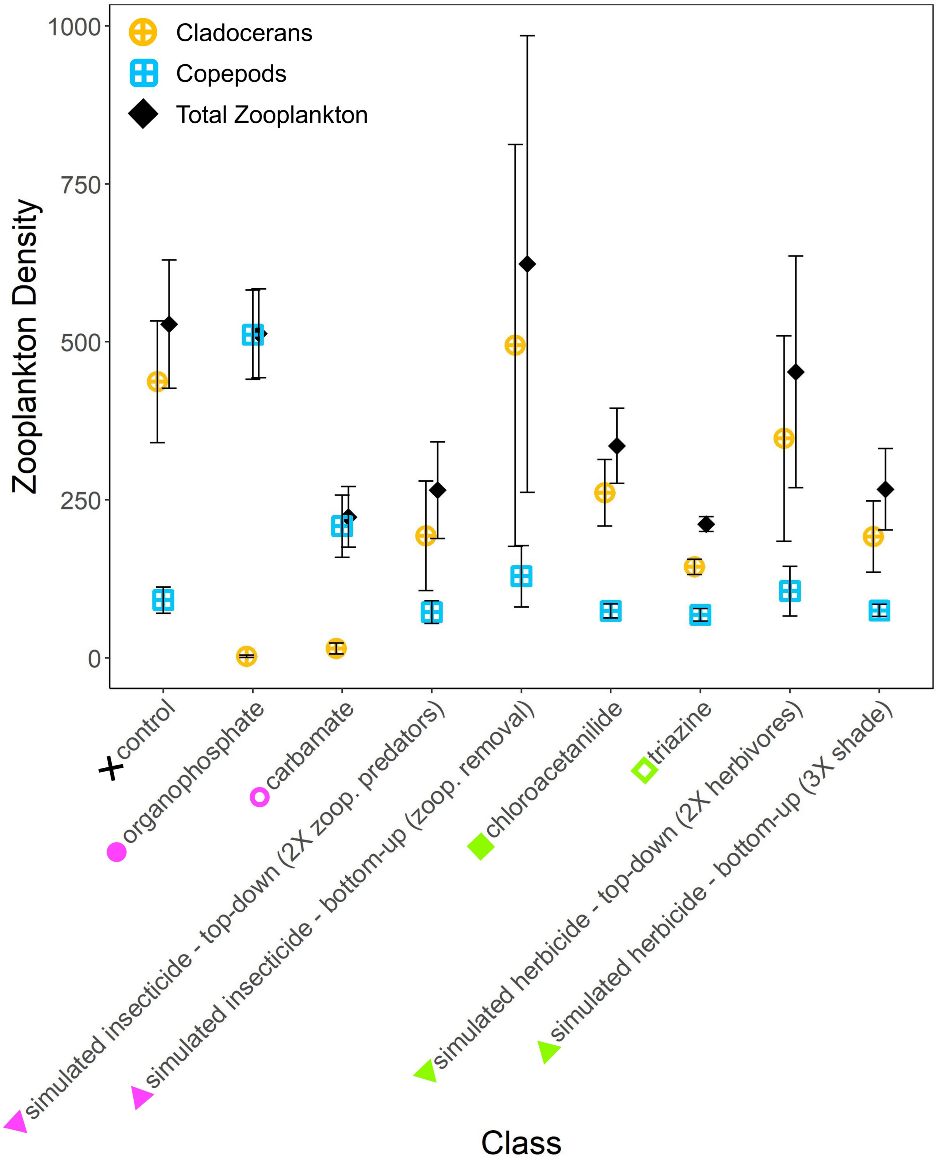


**Fig. S3 Zooplankton densities in response to experimental treatments.**
Densities of cladocerans, copepods, and total zooplankton in response to pesticide classes, simulated insecticides and herbicides, and the controls. The main impact of insecticides on zooplankton communities was a change in community composition with copepods becoming more abundant compared to cladocerans. In contrast, the main effect of herbicides on zooplankton was a decline in total abundance with no change in community composition; the relative amounts of cladocerans to copepods were comparable to the controls.

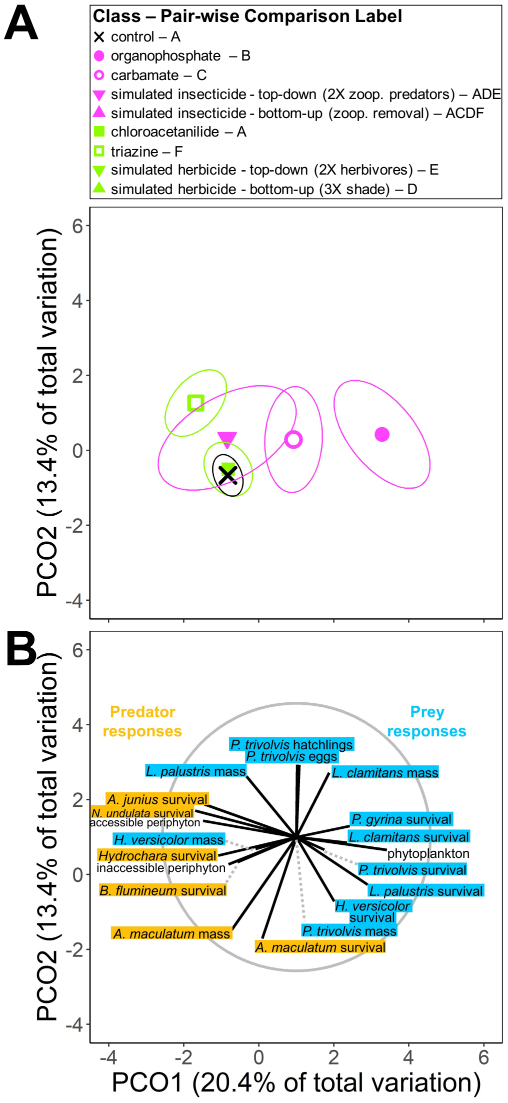


**Fig. S4 Principal coordinates analysis for multivariate community responses.**

**A)** Principal coordinates analysis plot of multivariate community-level responses showing differences in pesticide treatments by type, class, and individual pesticide. Individual points are centroids for the representative treatment with ellipses based on 95% confidence intervals calculated using a standard error. Simulated herbicides and insecticides that were different from corresponding pesticide treatments, including bottom-up simulated insecticide and top-down and bottom-up simulated herbicides, were not included in the plot for ease of viewing. Pair-wise comparison labels are given in the figure legend. Treatments sharing letters are not different from each other. **B)** Vector overlay of log-transformed and normalized community responses for corresponding principal coordinates analysis plot. Gray circle shows relative vector distance lengths; the gray circle corresponds to vector lengths that would have a correlation coefficient of one.

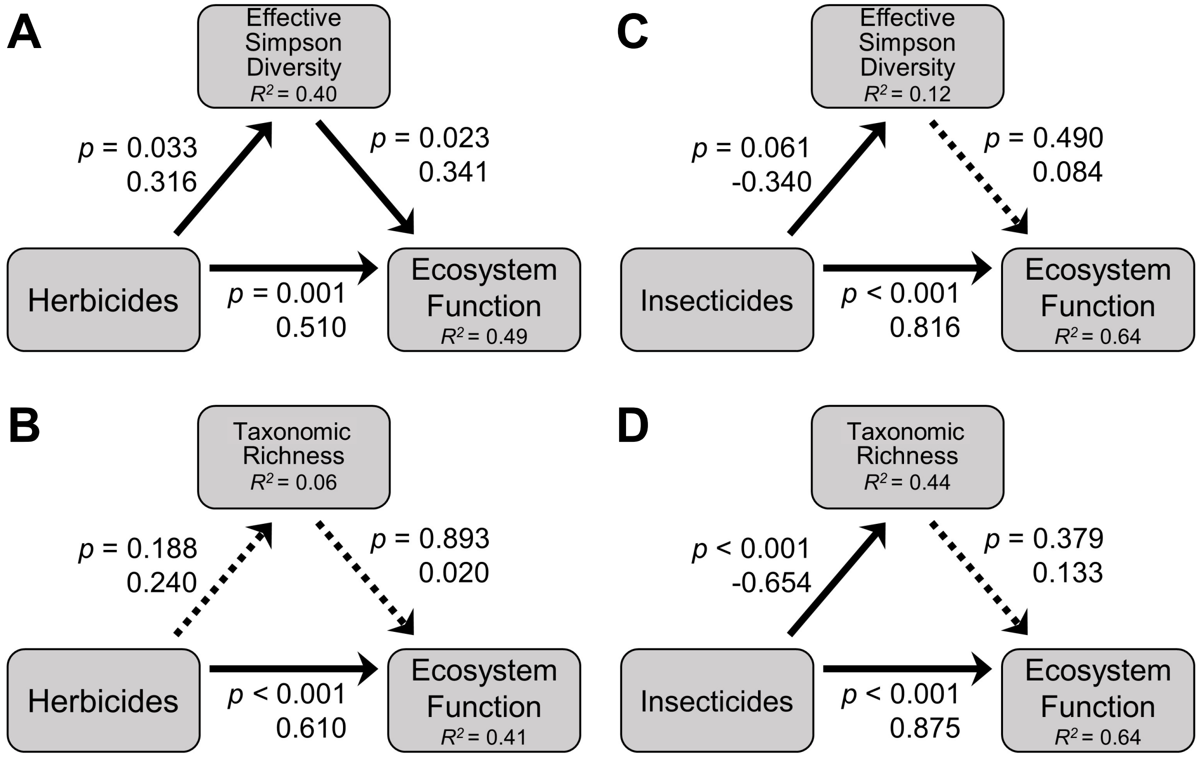


**Fig. S5 Relationships among pesticide types, diversity metrics, and ecosystem function.** Path analyses showing relationships among **A)** herbicides, Simpson diversity, and ecosystem function; **B)** herbicides, taxonomic richness, and ecosystem function; **C)** insecticides, Simpson diversity, and ecosystem function; and **D)** insecticides, taxonomic richness, and ecosystem function. For all path models, ecosystem function is the first axis from a principal coordinates analysis of the Euclidean resemblance matrix of log-transformed and normalized ecosystem responses including: pH and dissolved oxygen taken both at dawn and dusk, decomposition, and turbidity. Solid arrows are paths with $p < 0.07$, and dotted arrows are paths $p > 0.07$. Next to each path is the p-value and standardized coefficient. Next to each response is the conditional $R^2$.

**Table S1.**

PERMANOVA models evaluating the effects of pesticides on multivariate responses including ecosystem responses (pH and DO at dawn and dusk; decomposition; turbidity; periphyton; and phytoplankton), zooplankton densities (densities of six genera), tri-trophic community responses (survival of all non-zooplankton species; average mass of surviving amphibians and *H. trivolvis* snails; average eggs and hatchlings of surviving *P. trivolvis* snails; periphyton; and phytoplankton), and simplified tri-trophic community (combined responses of algae, herbivores, and predators). All models account for the influence of a spatial block. *P* values were generated by Monte Carlo sampling and those less than 0.05 are bolded. Variation explained, represented as a proportion, is the estimated component of variation for a given predictor relative to the model's total variation excluding block. So, variation explained accounts for the influence of the spatial block.

| Endpoints and Source of Variation | *df* | Pseudo *F* | *p* | Variation Explained |
|---|---|---|---|---|
| Ecosystem | | | | |
| Block | 3 | 2.142 | **0.013** | |
| Type | 1 | 21.247 | **0.0004** | 0.461 |
| Class(Type) | 2 | 1.346 | 0.224 | 0.073 |
| Pesticide(Class(Type)) | 8 | 1.838 | **0.004** | 0.146 |
| Residual | 33 | | | 0.319 |
| Zooplankton community | | | | |
| Block | 3 | 1.5395 | 0.124 | |
| Type | 1 | 9.6265 | **0.020** | 0.442 |
| Class(Type) | 2 | 4.551 | **0.004** | 0.188 |
| Pesticide(Class(Type)) | 8 | 1.8831 | **0.010** | 0.118 |
| Residual | 33 | | | 0.252 |
| Tri-trophic community | | | | |
| Block | 3 | 1.915 | **0.005** | |
| Type | 1 | 2.849 | **0.038** | 0.222 |
| Class(Type) | 2 | 1.806 | **0.034** | 0.154 |
| Pesticide(Class(Type)) | 8 | 2.111 | **0.0001** | 0.215 |
| Residual | 33 | | | 0.409 |
| Simplified tri-trophic community | | | | |
| Block | 3 | 2.697 | **0.012** | |
| Type | 1 | 4.271 | 0.087 | 0.291 |
| Class(Type) | 2 | 2.484 | 0.071 | 0.176 |
| Pesticide(Class(Type)) | 8 | 1.924 | **0.025** | 0.173 |
| Residual | 33 | | | 0.360 |

**Table S2.**

Model parameters of GENEEC version 2 used to generate environmentally relevant pesticide concentrations (Peak EEC [estimated environmental concentration]) used in the experiment.

| | Triazine herbicides | | | Chloroacetanilide herbicides | | | Carbamate insecticides | | | Organophosphate insecticides | | |
|---|---|---|---|---|---|---|---|---|---|---|---|---|
| | Atrazine | Propazine | Simazine | Acetochlor | Alachor | Metolachlor | Aldicarb | Carbaryl | Carbofuran | Chlorpyrifos | Malathion | Terbufos |
| Model Parameter Inputs | | | | | | | | | | | | |
| Trade name | Aatrex | Milocep | Princel 4L | Harness | Bullet | Dual II Magnum | Temik | Sevin 80S | Furadan | Dursban 50W | Fyfanon ULV | Counter 15G |
| Crop | Corn | Sorghum | Corn | Corn | Corn/ sorghum | Corn | Potatoes | Corn/ sorghum | Tobacco/ Barley | Turfgrass | Mosquito control | Corn |
| Application Rate (lbs a.i./acre) | 2 | 2 | 4.4 | 3 | 2.8125 | 2.3875 | 3 | 2 | 1.624 | 8 | 6 | 7.395 |
| Number of applications | 1 | 1 | 1 | 1 | 1 | 1 | 1 | 4 | 1 | 1 | 1 | 1 |
| Days between applications | - | - | - | - | - | - | - | 7 | - | - | - | - |
| $K_d$ | - | - | 1.96[a] | 3.03[a] | - | - | 0.053[a] | - | 1.23[a] | - | - | - |
| $K_{oc}$ | 100[a] | 65[d] | - | - | 170[a] | 200[a] | 30[a] | 300[a] | - | 6070[a] | 1248[b] | 500[a] |
| Soil half-life (d) | 300[b] | 231[e] | 100[c] | 84[a] | 49[c] | 56[c] | 72[a] | 21[a] | 120[a] | 30.5[b] | 6[c] | 5[b] |
| Wetted application? | No | No | No | No | No | No | No | No | No | No | No | No |
| Application method | Ground spray | Ground spray | Ground spray | Ground spray | Ground spray | Ground spray | Granular (2 inches) | Ground spray | Aerial | Ground spray | Ground spray | Granular (surface) |
| No spray zone (ft) | 0 | 0 | 0 | 0 | 0 | 0 | 0 | 0 | 0 | 0 | 0 | 0 |
| Solutbility (mg/L) | 33 | 8.5[b] | 5[a] | 223 | 242 | 530 | 6000[a] | 40[a] | 320[a] | 2[a] | 130[a] | 5 |
| Aquatic half-life (d) | 742[c] | 462[f] | 700[c] | 12[g] | 98 | - | 10[a] | 10[a] | 57[a] | - | - | 3.5[c] |
| Hydrolysis half-life (d) | - | - | - | - | - | 210 | - | - | - | 78[c] | 147[c] | - |
| Photolysis half-life (d) | 335[d] | - | - | - | - | 71 | 12[b] | 45 | 5[b] | 28[c] | - | - |
| Peak EEC (ppb) | 102 | 106 | 202 | 123 | 127 | 105 | 91 | 219 | 209 | 64 | 101 | 171 |

a Exotoxnet
b USDA
c Spectrum Laboratories
d USEPA fact sheet
e Pesticide Action Network
f Two times the soil half-life
g http://pmep.cce.cornell.edu/profiles/herb-growthreg/24-d-butylate/acetochlor/new-ai-acetochlor.html
